# A novel *PSMB8* isoform associated with multiple sclerosis lesions induces P-body formation

**DOI:** 10.1101/2024.02.26.582162

**Authors:** Benjamin C. Shaw, Jessica L. Williams

**Affiliations:** Department of Neurosciences, Lerner Research Institute, Cleveland Clinic, Cleveland, OH, USA; Brain Health Research Institute, Kent State University, Kent, OH, USA

**Keywords:** nonsense-mediated decay, neuroinflammation, alternative splicing

## Abstract

Multiple sclerosis (MS) is an inflammatory and demyelinating disease of the central nervous system (CNS). Current therapies primarily target the inflammatory component of the disease and are highly effective in early stages of MS while limited therapies have an effect in the more chronic progressive stages of MS where resident glia have a larger role. MS lesions tend to be inflammatory even after the initial peripheral immune cell invasion has subsided and this inflammation is known to cause alternative splicing events. We used qPCR of normal-appearing white matter and white matter lesions from postmortem MS tissue, *in vitro* studies, and immunostaining in MS tissue to investigate the alternative splicing of one gene known to be important during recovery in an animal model of MS, *PSMB8*. We found a novel, intron-retained isoform which has not been annotated, upregulated specifically in MS patient white matter lesions. We found that this novel isoform activates the nonsense-mediated decay pathway in primary human astrocytes, the most populous glial cell in the CNS, and is then degraded. Overexpression of this isoform in astrocytes leads to an increased number of processing bodies *in vitro*, the primary site of mRNA decay. Finally, we demonstrated that MS white matter lesions have a higher burden of processing bodies compared to normal-appearing white matter, predominantly in GFAP-positive astrocytes. The increase in alternative splicing of the *PSMB8* gene, the stress that this alternative splicing causes, and the observation that processing bodies are increased in white matter lesions suggests that the lesion microenvironment may lead to increased alternative splicing of many genes. This alternative splicing may blunt the protective or reparative responses of resident glia in and around white matter lesions in MS patients.

## Introduction

Multiple sclerosis (MS) is a chronic inflammatory, neurodegenerative disease of the central nervous system (CNS) (Compston and Coles, 2008). Histologically, MS lesions are characterized by peripheral immune cell infiltration, demyelination, and glial cell activation (Trapp and Nave, 2008). The most common clinical subtype of MS is relapsing remitting MS (RRMS). Patients with RRMS exhibit periods of exacerbated symptoms such as paresthesia, vision deficits, and muscle weakness, followed by periods of partial to complete recovery (Compston and Coles, 2008). Most of these patients will develop secondary progressive MS (SPMS), and patients in this stage of MS exhibit markedly less recovery between relapses (Hurwitz, 2009). A small proportion of MS patients will initially present with a progressive phenotype, termed primary progressive MS (PPMS), characterized by constant disability accumulation with no clear remission (Confavreux et al., 2000, Thompson et al., 1991). At any stage of the disease, patients can experience disability progression independent of relapse activity (PIRA), which is the most common form of disability accumulation across all traditional MS subtypes (Müller et al., 2023). Patients with RRMS, typically characterized by a significant inflammatory component, respond well to current disease-modifying therapies, such as ocrelizumab, fingolimod, and glatiramer acetate, which are immunomodulatory (Wingerchuk and Carter, 2014). In progressive MS, however, these immune cell-targeting drugs have limited efficacy indicating that disability is perpetuated by CNS resident cells (Miller and Leary, 2007, Feinstein et al., 2015).

Astrocytes are the most abundant glial cell in the CNS and are responsible for the formation of the glial scar in MS lesions (Brosnan and Raine, 2013). Traditionally thought of as quiescent support cells, recent studies have highlighted their importance in both protection from and induction of neuroinflammation (Lee et al., 2023). Astrocytes are exquisitely responsive to immune signaling molecules, including interferons, commonly found in MS lesions (Daniels et al., 2017, Williams et al., 2020). Specifically, interferon-γ (IFNγ) upregulates the immunoproteasome within astrocytes as a protective mechanism in a murine model of MS, experimental autoimmune encephalomyelitis (EAE) (Smith et al., 2020). The immunoproteasome consists of multiple subunits—LMP7 (*PSMB8*), LMP2 (*PSMB9*), and MECL-1 (*PSMB10*)—which are induced by interferons and assemble sequentially (Griffin et al., 1998, Heink et al., 2005, Orre et al., 2013). Once assembled, the immunoproteasome is responsible for the proteolysis of polyubiquitinated and oxidatively damaged proteins (Ebstein et al., 2012).

The immunoproteasome is upregulated in MS lesions, especially in astrocytes, and inhibition of the immunoproteasome exacerbates EAE (Smith et al., 2020). Counterintuitively, activity of the immunoproteasome is decreased in MS lesions despite its protein-level upregulation (Zheng and Bizzozero, 2011). Alternative splicing can lead to a loss of function for genes critical in regulating other diseases, such as *TREM2* in Alzheimer’s disease (Shaw et al., 2022, Kiianitsa et al., 2021). Given the sequential assembly of the immunoproteasome (Griffin et al., 1998), wherein LMP7 is the first subunit incorporated followed by LMP2 and then MECL-1, disruption of either LMP7 protein or its transcript, *PSMB8*, may lead to an incomplete or inefficient induction. Poor induction of the immunoproteasome may therefore lead to further decreases in immunoproteasome activity and thus increased lesion activity. Further, inflammation induces alternative splicing of many mRNAs and alternative splicing frequently leads to premature stop codon insertion which activates the nonsense-mediated decay (NMD) pathway and induces processing bodies (P-bodies) (Janssen et al., 2020, Robinson et al., 2021). Thus, the inflammation which occurs in an MS lesion may lead to alternative splicing of critical damage control mechanisms limiting the potential recovery after CNS insult. To date, however, no studies have examined alternative splicing in CNS tissue at patient lesion sites during MS.

To address this, we focused our attention on the immunoproteasome gene *PSMB8*. This gene has a pair of mutually exclusive exons, exon 1A and exon 1B. Importantly, only the protein which results from translation of exon 1B-containing *PSMB8* can efficiently incorporate into the mature immunoproteasome (Griffin et al., 1998). While previous reports have found that inflammation can drive alternative first exon usage (Robinson et al., 2021), we found no difference in the proportion of exon 1A versus exon 1B usage in MS white matter lesions (WML) compared to normal-appearing white matter (NAWM). We did, however, uncover a previously unreported and unannotated transcript of *PSMB8* which retains intron 2 and is upregulated in WMLs. Using primary human cortical astrocytes, we found that this intron 2 retained *PSMB8* transcript (*i2R-PSMB8*) activates the NMD pathway and is then degraded. Further, overexpression of *i2R-PSMB8* increases the number of intracellular P-bodies *in vitro*. Finally, we observed a substantial increase in overall P-body burden within astrocytes among other cells within MS lesions, potentially indicating a dysregulation in splicing of many genes within the MS WML.

## Materials and Methods

### Human tissue, RNA isolation, and cDNA conversion

MS brain tissues were obtained from The Cleveland Clinic according to the established rapid autopsy protocol approved by the Cleveland Clinic Institutional Review Board (Chomyk et al., 2017). Patients or their relatives provided informed consent in the form of an advanced directive as part of an institutional review board-approved protocol. Tissues originating from MS cases used for qRT-PCR and immunohistochemistry are described in Table 1. Tissues were obtained, lesion types characterized, and RNA from human cerebral tissue was isolated and converted to cDNA as previously described (Dutta et al., 2006). Paired WML and NAWM were used for qRT-PCR analysis.

**Table 1.** Patient demographics.

| Patient ID | Sample Type | Age (yrs) | Sex | MS Type | EDSS | Postmortem Interval (hrs) | Disease Duration (Yr) |
| --- | --- | --- | --- | --- | --- | --- | --- |
| 31 | NAWM, WML | 60 | F | Secondary Progressive MS | 9 | 7.83 | 29.5 |
| 82 | NAWM, WML | 51 | F | Primary Progressive MS | 7.5 | 6.92 | 14.9 |
| 88 | NAWM, WML | 57 | M | Secondary Progressive MS | 6.5 | 9.67 | 27.5 |
| 144 | NAWM, WML | 38 | M | Secondary Progressive MS | 7 | 12 | 17.9 |
| 167 | NAWM, WML | 59 | F | Secondary Progressive MS | 7.5 | 9.5 | NA |
| 170 | NAWM, WML | 48 | F | Relapsing Remitting MS | 7.5 | 10 | NA |

### End-point and qRT-PCR assays

Within patient WML and NAWM cDNA samples were amplified using a forward primer against exon 1B (5’-GCTCGGACCCAGGACACTAC-3’) and a reverse primer against exon 6 (5’-GAGAACACGCAGAAGATGCAC-3’) with AccuStart II PCR Supermix (QuantaBio) for an end-point PCR assay. Samples were amplified using an initial 1 min denaturation at 95°C, followed by cycles of 15 s at 95°C, 15 s at 57.3°C, and 2 min at 72°C for 40 cycles, followed by a 10 min final extension at 72°C before resolving on a 5% polyacrylamide tris-borate-EDTA (TBE) gel (Bio-Rad) at 35 V. Gel bands were excised, reamplified, and purified using a post-PCR cleanup kit (New England Biolabs) per manufacturer’s instructions. Purified DNA was sequenced commercially (Azenta) to identify excised bands. For qRT-PCR assays, cDNA was amplified using a forward primer in exon 1B (5’-GCTCGGACCCAGGACACTAC-3’) and a reverse primer in exon 2 (5’-GTTCCTTTCTCCGTCCCCAC-3’) to quantify total exon 1B-containing *PSMB8*. Intron 2 retention was quantified with a forward primer in intron 2 (5’-CCCTTCTCTCCCAAAGCTCC-3’) and reverse primer in exon 3 (5’-GTGCCAAGCAGGTAAGGGTT-3’). *GAPDH* was used as a housekeeping gene and quantified (5’-GAAGGTGAAGGTCGGAGTC-3’ and 5’-GAAGATGGTGATGGGATTTC-3’). All qRT-PCR assays used PowerSYBR master mix (Applied Biosystems), and assays were performed according to manufacturer’s instructions in duplicate and quantified using the double-delta Ct method.

### Cell culture and transfections

Primary human cortical astrocytes (ScienCell) were obtained commercially and expanded in astrocyte media (ScienCell). Cells were cultured at 37°C in 5% CO_2_ and split at approximately 80% confluency until passage 3. Upon reaching passage 3, cells were frozen using Recovery Cell Freezing Media (Gibco). Cells were thawed prior to experiments and transfected during passage 4. Vectors encoding either intron 2 retained *PSMB8* (*i2R-PSMB8*) or the canonical full-length transcript (*FL-PSMB8*) in pcDNA3.1 were obtained from GenScript, outgrown in DH5α *E. coli* (New England Biolabs), and midi-prepped using a Plasmid Plus Midi kit (Qiagen) per manufacturer’s instructions. Vector maps are provided as Supplemental Figures 1-2. Vectors were transfected using a Lonza Amaxa Nucleofector 4 device, with kit P3 and protocol DR114, at a ratio of 6 μg DNA per 10^6^ cells. This resulted in a 50-55% transfection efficiency (**Supplemental Figure 3**).

### Protein isolation and western blotting

Cells were cultured in 6-well plates after transfection at 80% confluency and incubated for 24 h post-transfection. Cells were then washed three times with PBS and lysed with IP lysis buffer (25 mM Tris-HCl, 150 mM NaCl, 1 mM EDTA, 1% Nonidet P-40, 5% glycerol) with 1% protease-phosphatase inhibitor (Pierce). Debris was pelleted at 15,000 RCF for 15 min at 4°C and protein concentration determined using a BCA assay (Invitrogen). Protein (50 μg per sample) was reduced using β-mercaptoethanol, denatured using SDS loading buffer (Bio-Rad) and heat denatured at 95°C for 10 min before loading onto a 4-15% Tris-glycine gel (Bio-Rad) and electrophoresed at 125 V. Gels were transferred to PVDF membranes using a Turbo transfer system (Bio-Rad). Membranes were blocked with 3% bovine serum albumin in Tris-buffered saline with 0.05% Tween-20 (TBS-T) for 1 h at room temperature and blotted with an antibody against phosphorylated upstream frameshift protein 1 (phospho-UPF1) (pUPF1; 1:1000, Cell Signaling Technologies) overnight at 4°C. Membranes were then washed three times with TBS-T prior to incubating with a goat anti-rabbit horseradish peroxidase (HRP) conjugated antibody (Invitrogen). Chemiluminescence was detected using an enhanced chemiluminescence kit (Bio-Rad) and a ChemiDoc MP system (Bio-Rad). After imaging, membranes were stripped twice for 10 min each using stripping buffer (200 mM glycine, 3.5 mM SDS, 1% Tween-20, pH 2.2), blocked with 5% non-fat milk, and blotted overnight at 4°C with an antibody against total UPF1 (1:1000, Cell Signaling Technologies). Membranes were again washed three times with TBS-T prior to incubating with a goat anti-rabbit HRP conjugated antibody (Invitrogen), and chemiluminescence detected as before.

### *PSMB8* isoform stability

Stability of mRNA isoforms was assayed as previously described (Zajac et al., 2023). Briefly, primary human cortical astrocytes (ScienCell) were cultured to passage 4 and stimulated with 50 μg/mL cycloheximide (CHX; Invitrogen) or dimethylsulfoxide (DMSO) vehicle control for 0, 1, 4, or 8 h. The assay was performed in triplicate per time point and treatment. At the designated time point, cells were lysed and RNA was collected using an RNeasy isolation kit (Qiagen) per manufacturer’s instructions. Resultant RNA was converted to cDNA using 500 ng total RNA and a high-capacity reverse transcription kit (Applied Biosystems, 100025924) with final concentrations of reagents as follows: 1.75 mM MgCl_2_ (Invitrogen), 2.00 mM dNTPs (Applied Biosystems), 2.50 μM random hexamers (Invitrogen), 2.50 μM oligo(dT) (Invitrogen), 1 U/μL RNAse inhibitor (Applied Biosystems), 5 mM dithioerythritol (Invitrogen), 2.50 U/μL reverse transcriptase (Applied Biosystems). The 20 μL reactions were heated to 37°C for 30 min then heat-inactivated at 95°C for 5 min before storage at -20°C until qPCR was performed.

### Immunocytochemistry and imaging of transfected cells

Transfected cells were washed once with PBS then fixed in 4% paraformaldehyde for 30 min. Cells were blocked and permeabilized in 10% normal goat serum and 0.1% Triton-X100 in PBS for 1 h. Cells were then stained with an antibody against enhancer of mRNA decapping 4 (EDC4) (1:200, Abcam) at room temperature for 1 h before washing three times with PBS. Secondary antibody (Invitrogen) was then applied for 1 h, slides washed three times with PBS, and incubated with DAPI solution (Invitrogen) to stain nuclei. Slides were coverslipped with Prolong Glass mounting media (Invitrogen) and stored at -20°C until imaged using a Zeiss LSM800 confocal microscope. ImageJ was used to automate EDC4^+^ puncta counts in GFP+ cells to remove bias.

### Immunohistochemistry of human MS tissue

Paraformaldehyde-fixed tissue sections (30 μm thick) were obtained from the Cleveland Clinic rapid autopsy program. Antigen retrieval was performed using 10 μM citrate buffer and boiling briefly. Free-floating sections were permeabilized with 2% Triton-X100 in PBS for 30 min then blocked with 5% normal goat serum and 0.3% Triton-X100 for 1 h. Sections were then incubated in primary antibody against major histocompatibility complex class II (MHCII; 1:50, Abcam), myelin basic protein (MBP; 1:200, Abcam), glial fibrillary acidic protein (GFAP; 1:50, Invitrogen), and EDC4 (1:40, Abcam) for three days. Sections were then washed in PBS with 0.3% Triton-X100 three times and incubated with secondary antibodies (Invitrogen) conjugated to AlexaFluor 405, 488, 594, or 647 for 1 h at room temperature. Slides were then washed three times and treated with TrueBlack lipofuscin quencher (Biotium) per manufacturer’s instructions. Sections were mounted using Prolong Glass (Invitrogen) and stored at -20°C until imaged using a Zeiss LSM800 confocal microscope. Cell bodies filled with EDC4 were counted by a blinded observer.

### Statistical analyses

Quantitative PCR data using the double-delta Ct method were analyzed using a one-sample *t* test. Western blot, puncta count, and cell body count data were analyzed using a two-sample *t* test.

## Results

### Discovery of an alternative splice variant in *PSMB8*

To address whether any novel splice variants of the functional exon 1B-containing *PSMB8* exist, we amplified cDNA from MS patients derived from either NAWM or WMLs with a forward primer targeting exon 1B and a reverse primer targeting exon 6. After amplification and polyacrylamide gel electrophoresis, the canonical isoform of *PSMB8* was highly expressed in both NAWM and WMLs. Additionally, a higher molecular weight band indicating a larger transcript was apparent in WMLs (**Figure 1A**). This band was excised, reamplified, and sequenced. Sequencing revealed that this novel isoform retains intron 2, leading to an additional 158 bp in the resultant mRNA, consistent with the molecular weight shift observed in WMLs (**Figure 1A**). This novel isoform is hereafter referred to as intron 2 retained or *i2R-PSMB8*. Quantitative PCR of the same MS patient cDNA demonstrated that exon 1B usage, relative to *GAPDH*, is significantly increased (**Figure 1B**; *p* = 0.0005) in WMLs compared to NAWM consistent with previous reports (Smith et al., 2020). Additionally, qPCR using primers targeting intron 2 and exon 3 revealed that intron 2 retention was significantly higher in WMLs compared to NAWM relative to total exon 1B expression (**Figure 1C**; *p* < 0.0001). While the band we identified as *i2R-PSMB8* in Figure 1A appears to represent a small amount of the total *PSMB8*, we hypothesize that this isoform is predicted to undergo NMD and is likely highly underrepresented on the gel image. These data indicate that in the lesion microenvironment, *PSMB8* splicing is dysregulated and may contribute to increased cellular stress.

**Figure 1.**
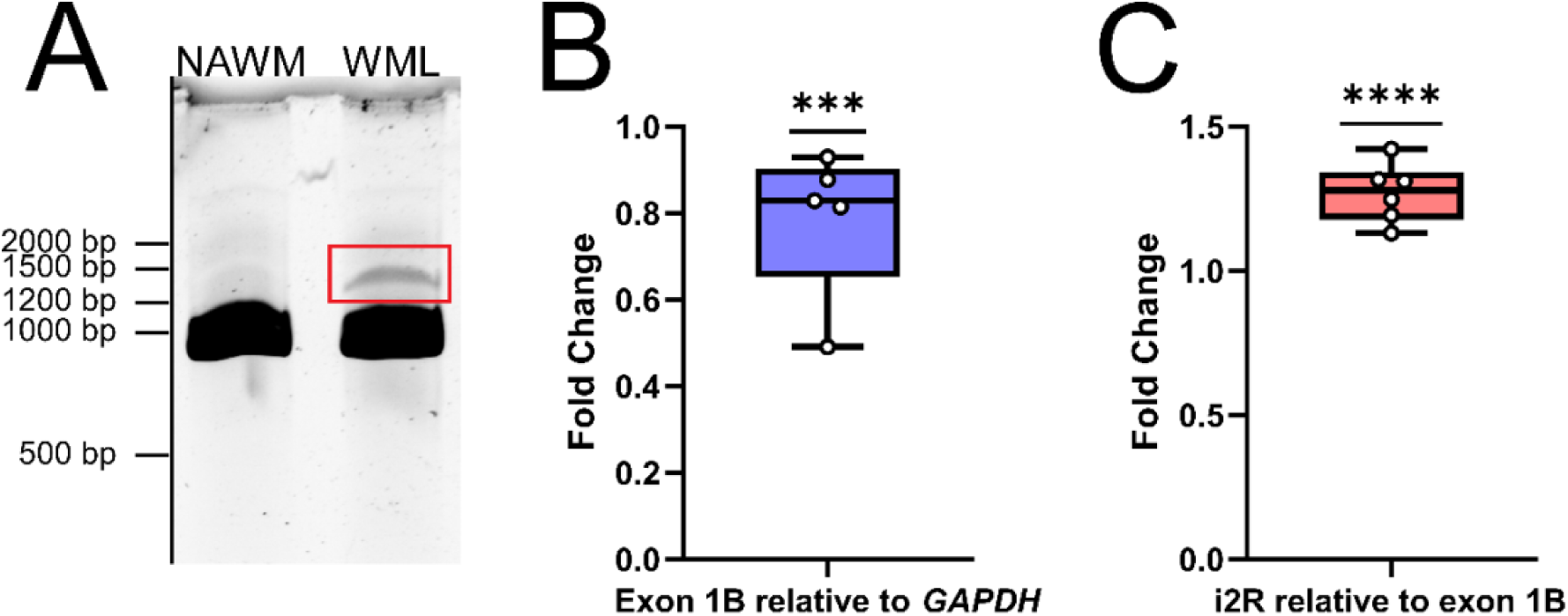
*PSMB8* undergoes alternative splicing in the MS WML. **(A)** Representative gel image of cDNA amplified with primers corresponding to exon 1B and exon 6 of *PSMB8*. cDNA was amplified from NAWM (left lane) or WML (right lane) to visualize the canonical PSMB8 isoform (918 bp) and a higher molecular weight isoform (1076 bp, red box). **(B)** qPCR for total exon 1B was performed on cDNA from NAWM and WMLs. *PSMB8* expression was then normalized to *GAPDH*. One sample was excluded due to lack of amplification in *GAPDH*. **(C)** *i2R-PSMB8* was quantified and normalized to total exon 1B usage. Data representative of two independent experiments. Significance was determined using a one-sample t-test. ** *p* < 0.01. *** *p* < 0.001.

### *i2R-PSMB8* enhances P-body formation in astrocytes

Since the *i2R-PSMB8* isoform has an additional 158 bp inserted after exon 2, there is a frameshift and resultant premature stop codon in exon 3. Given that *PSMB8* has six exons, we hypothesized that the *i2R-PSMB8* isoform would undergo NMD. P-bodies are the primary sites of mRNA decay after being targeted through the NMD pathway (Teixiera et al., 2005). Increases in NMD are correlated with increased size and number of P-bodies (Teixiera et al., 2005, Di Stefano et al., 2019, Chuang et al., 2013, Chang and Tarn, 2009), which can be quantified using EDC4, a critical component of P-body formation. To test this hypothesis, we cloned the *i2R-PSMB8* and *FL-PSMB8* transcripts and transfected these vectors into human primary cortical astrocytes in glass chamber slides and co-transfected with a vector encoding green fluorescent protein (GFP) to mark transfected cells (**Figure 2A-B**). At 24 h post-transfection, cells were fixed and labeled for EDC4. Notably, the number of EDC4^+^ puncta within GFP^+^ cells was significantly higher in the *i2R-PSMB8* transfected astrocytes compared to those transfected with *FL-PSMB8* (**Figure 2C**; *p* = 0.0003). This indicates the *i2R-PSMB8* transcript may lead to increased NMD. To test this, we again transfected primary human cortical astrocytes with the *i2R-PSMB8* or *FL-PSMB8* vectors and quantified the phosphorylation of UPF1, an NMD initiating event (Kashima et al., 2006). At 24 h post-transfection, we assessed the ratio of pUPF1 to total UPF1 via western blot (**Figure 2D**) and found that there was significantly enhanced activation of UPF1 in cells transfected with *i2R-PSMB8* compared to *FL-PSMB8* (**Figure 2E**; *p* = 0.0196).

**Figure 2.**
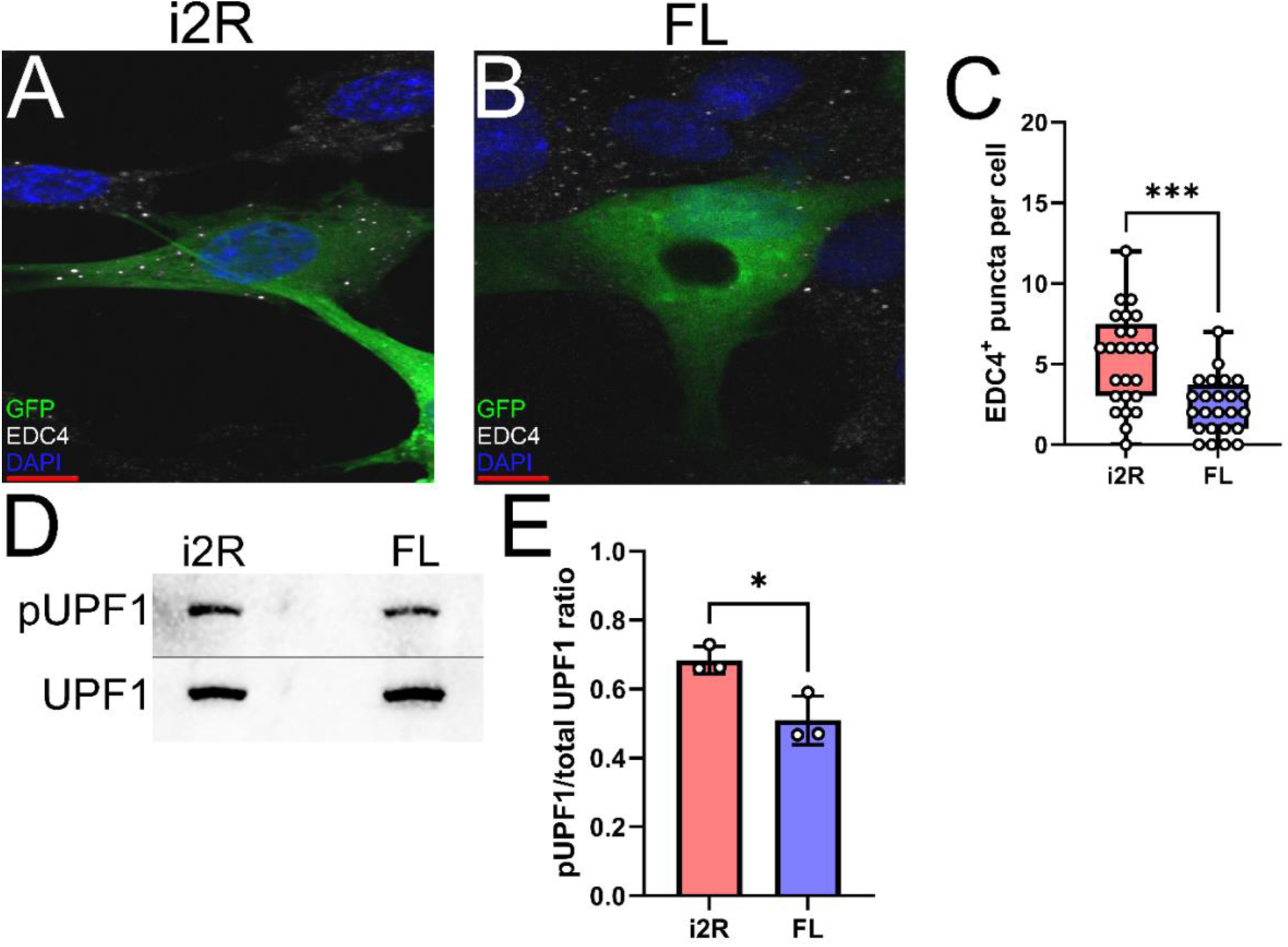
*i2R-PSMB8* induces P-body formation and leads to phosphorylation of UPF1. Human primary cortical astrocytes were transfected with vectors encoding either **(A)** intron 2 retained (i2R) or **(B)** full-length (FL) *PSMB8* and co-transfected with a vector encoding green fluorescent protein (GFP) to identify positively transfected cells. **(C)** GFP^+^ cells were identified and the automated counting of the number of EDC4^+^ puncta (white) was performed using ImageJ. Scale bars (red) are 10 μm. Data representative of two independent experiments. Each point represents an individual transfected GFP^+^ cell. **(D)** Representative western blot images of phospho-UPF1 (top) and total UPF1 (bottom) of lysates from primary human cortical astrocytes transfected with vectors encoding either intron 2 retained (i2R) or full-length (FL) *PSMB8*. **(E)** The ratio of pUPF1 to total UPF1 was quantified. Data are from three independent experiments. Significance was determined by unpaired *t* test. * *p* < 0.05. *** *p* < 0.001.

### The *i2R-PSMB8* transcript undergoes NMD in astrocytes

The NMD pathway is dependent on first-pass translation (Behm-Ansmant and Izaurralde, 2006), and inhibiting translation through treatment with cycloheximide causes transcripts typically targeted for NMD to become more stable (Zajac et al., 2023). To directly test whether the *i2R-PSMB8* transcript undergoes NMD, we treated primary human cortical astrocytes with either cycloheximide (50 μg/mL) or DMSO as a vehicle control and incubated for up to 8 h. Total RNA was isolated at 0, 1, 4, and 8 h post-treatment and converted to cDNA for qPCR. Subsequent qPCR revealed a time-and drug-dependent interaction (**Figure 3**; *F_3,16_ =* 18.09*, p* < 0.0001), indicative of an increase in *i2R-PSMB8* transcript in the cycloheximide-treated, but not vehicle-treated astrocytes, suggesting that the *i2R-PSMB8* transcript undergoes NMD.

**Figure 3.**
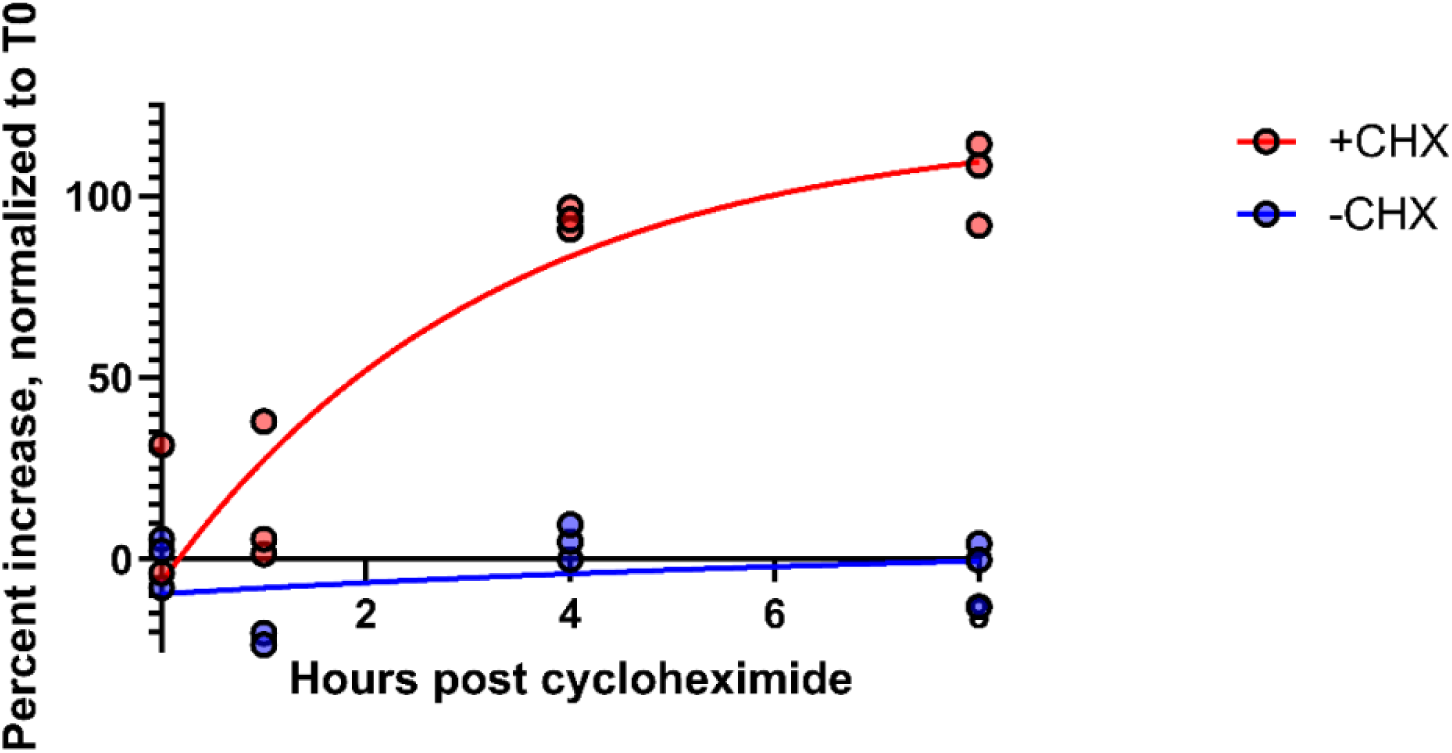
*i2R-PSMB8* undergoes NMD in astrocytes. qPCR measurements of intron 2 retained (i2R) transcript after the addition of cycloheximide (+CHX) or dimethylsulfoxide (DMSO) as a vehicle control (-CHX). Data are normalized to time 0 and are representative of two independent experiments. Significance was determined by two-way ANOVA. *F_3,16_ =* 18.09*, p* < 0.0001.

### P-body load is increased in postmortem MS lesions

Since MS lesions preferentially contain higher levels of *i2R-PSMB8* transcript compared to NAWM (**Figure 1**) and that this transcript leads to enhanced P-body formation and NMD (**Figure 2-3**), we next asked whether MS lesions also displayed an increase in P-body formation in astrocytes. To address this question, we obtained postmortem MS tissue sections and labeled them for MBP and MHCII to determine lesion type along with GFAP and EDC4 to determine P-body burden within astrocytes. We chose to analyze chronic active WMLs due to their active lesion border that is dense with reactive astrocytes {Zeinstra, 2003 #1256}. In confirmed chronic active lesion borders of MS patients (**Figure 4Ai-iii**), we observed striking densities of EDC4^+^ structures (**Figure 4A**), which appeared much larger than the P-bodies observed in our *in vitro* studies (**Figure 2A-B**). By contrast, much smaller EDC4^+^ structures were observed in MS patient NAWM (**Figure 4B**), which was distal to lesions and where MBP was apparently intact (**Figure 4Bi-iii**). Using 3D image rendering, we found that the large EDC4^+^ structures were frequently, but not exclusively, found within GFAP^+^ astrocytes (**Figure 4C**). Quantifying the number of these EDC4-laden astrocytes and normalizing to the number of GFAP^+^ cell bodies per high powered field, we observed an increasing trend in the proportion of EDC4-laden astrocytes in the chronic active lesion borders compared to in NAWM (**Figure 4D**; *p* = 0.0673). These data indicate that the astrocytes, among other cells, of the chronic active lesion border undergo significant alternative splicing events, which are likely to the detriment of proper cellular function.

**Figure 4.**
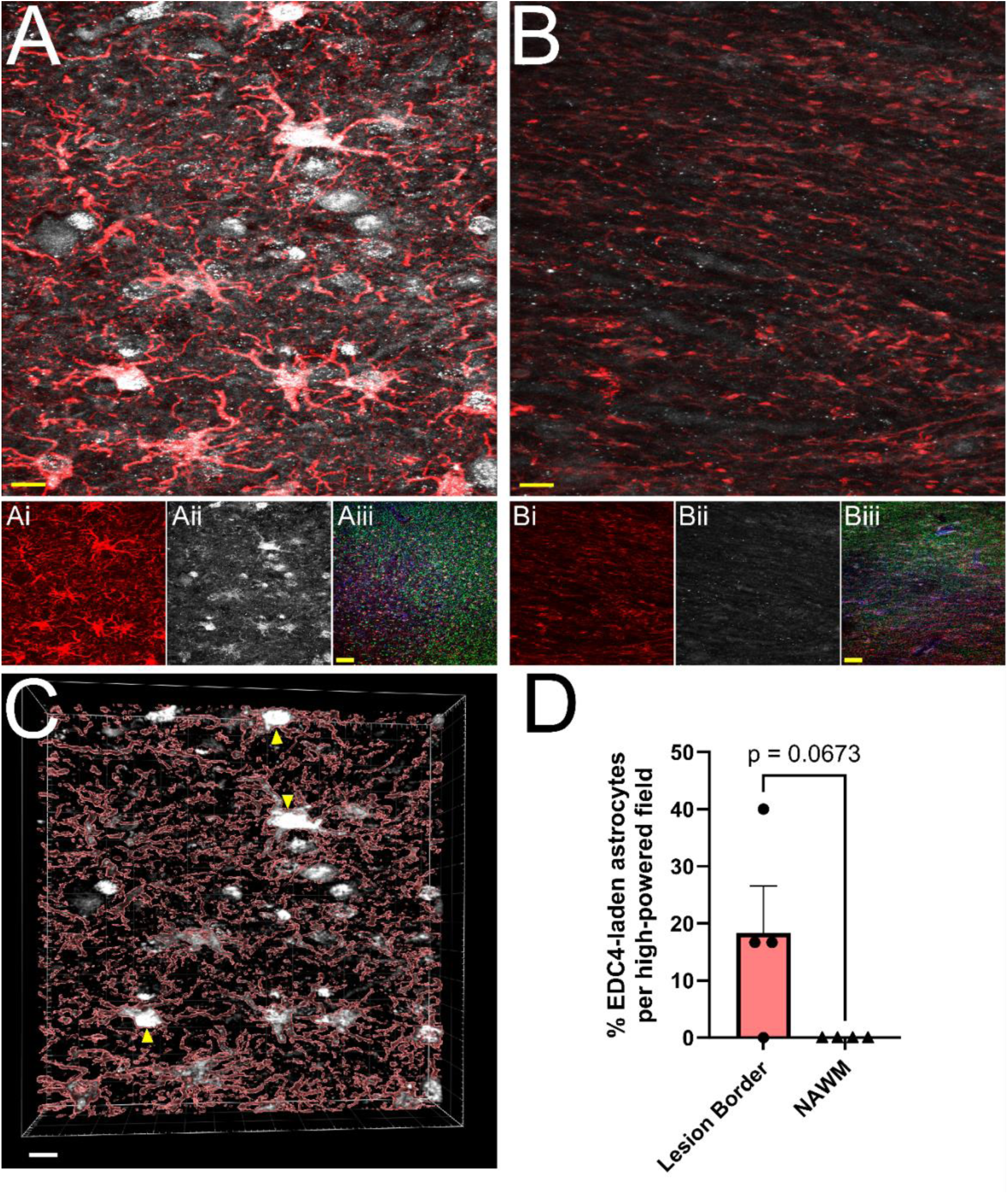
MS WMLs exhibit substantial EDC4 condensation. Representative merged confocal images of either **(A)** white matter lesion or **(B)** normal-appearing white matter stained for GFAP (red) and EDC4 (white). **(Ai-ii)** Single channel images of the representative chronic active lesion from MS patient 88. **(Aiii)** A representative merged 10X confocal image of the chronic active lesion characterization labeled for MHCII (Blue), MBP (Green), GFAP (Red), and EDC4 (White). **(Bi-ii)** Single channel images of the representative chronic active lesion from MS patient 167. **(Biii)** A representative merged 10X confocal image of tissue from patient 167 labeled for MHCII (Blue), MBP (Green), GFAP (Red), and EDC4 (White). Scale bars (yellow) in **(A)** and **(B)** are 10 μm. Scale bars (yellow) in **(Aiii)** and **(Biii)** are 100 μm. **(C)** 3D volumetric reconstruction of **(A)** showing buildup of EDC4 (white) in astrocytes (red; GFAP). Yellow arrowheads indicate EDC4-laden astrocytes. Scale bar is 10 μm. **(D)** Quantification of percent of astrocyte cell bodies laden with EDC4 per high powered field. Statistics by two-sample *t* test.

## Discussion

Multiple studies have shown a potential role for alternative splicing in neurodegenerative disorders (Shaw et al., 2022, De Rossi et al., 2016, Paraboschi et al., 2014, Putscher et al., 2022). In this study, we identified a novel *PSMB8* splice variant preferentially upregulated in demyelinated WMLs (**Figure 1A**), the primary pathology during MS, and corroborate previous work (Smith et al., 2020) demonstrating that *PSMB8* is upregulated in WMLs of MS patients (**Figure 1B**). Further, our work demonstrates that this novel splice variant undergoes NMD and leads to an increase in P-body formation *in vitro* (**Figure 2-3**). This preferential increase in a novel splice variant which is degraded in the WML led us to hypothesize that splicing may be globally dysregulated within the WMLs during MS. Indeed, we also found that large EDC4^+^ structures exist within cells of the WML, especially in astrocytes (**Figure 4**). These structures appeared to occupy a substantial volume of the GFAP^+^ cell bodies. These results suggest that in the lesion microenvironment, splicing is dysregulated within astrocytes which may further compound cellular stress.

Specifically, regarding the *i2R-PSMB8* isoform, this alternative splicing event may reduce the capacity for full induction of the immunoproteasome. While we found that total *PSMB8* is upregulated in the WML environment, a portion of this upregulation is nonfunctional as the alternative splicing which gives rise to *i2R-PSMB8* does not encode a functional protein (**Figure 1C**). The protein encoded by *PSMB8*, LMP7, is required for immunoproteasome assembly (Griffin et al., 1998). Loss of function of the immunoproteasome is detrimental in animal models of MS, such as EAE (Smith et al., 2020). Thus, this nonfunctional *i2R-PSMB8* isoform may represent a partial loss of induction of functional *PSMB8*. We note that while the *i2R-PSMB8* isoform appears to be a small portion of total in Figure 1A, this is actually an underestimation of the total amount made as we have now shown that it is substantially degraded through NMD. Whether the expression level of this isoform present is enough to reduce the functional LMP7 below optimal levels remains unknown; however, the LMP7 protein concentration does closely correlate well (*ρ* = 0.91) with the *PSMB8* mRNA counts, indicating a disruption in mRNA would likely have effects on functional protein (Jiang et al., 2020).

Beyond this specific isoform, though, it appears splicing as a whole may be dysregulated in the lesion microenvironment given the drastic increase in EDC4 labeling in MS lesions, as a single isoform is not likely to have such a profound impact. Proinflammatory cytokines induce alternative splicing in many genes *in vitro*, including in primary astrocytes derived from human MS patients (Werkman et al., 2020). Additionally, in animal models of MS, splicing is dysregulated in lesions and this aberrant splicing correlates with interleukin-1β expression (Marchese et al., 2021, Adinolfi et al., 2023). Additionally, the microglial splicing factor QKI-5 is downregulated in brain WMLs of MS patients (Lee et al., 2020). We show here that EDC4, a marker for NMD and a proxy for aberrant splicing, is highly upregulated in MS WMLs, frequently occurring in GFAP^+^ astrocytes, though other cells also show high levels of EDC4 expression. This most frequently occurred in the chronic active lesion border, where inflammation is thought to persist. This corresponds with preclinical observations correlating alternative splicing with inflammation (Marchese et al., 2021, Adinolfi et al., 2023).

It is possible that in MS lesions, the immunoproteasome is one of possibly many responses to inflammation that could be dampened due to alternative splicing. We speculate that this increased alternative splicing in the lesion microenvironment may contribute to lesion progression, limiting the ability of resident glia to repair damage. One limitation of our study is that we cannot determine whether the specific *i2R-PSMB8* transcript alone is sufficient to cause the substantial increase in EDC4 condensation observed in the MS lesion. We hypothesize that this one transcript alone is insufficient; however, and the more likely explanation is that many genes are undergoing detrimental alternative splicing in the WMLs to give rise to this EDC4-laden phenotype. Further, the observed increases in P-body formation may result in increased cell stress and death, possibly contributing to the slowly expanding smoldering lesions frequently found in progressive MS. Identifying specific splicing factors affected in MS lesions could lead to a viable therapeutic strategy.

## Ethics Statement

Brain tissues from MS patients (**Table 1**) were collected according to the established rapid autopsy protocol approved by the Cleveland Clinic Institutional Review Board.

## Acknowledgements

We thank Kaitlin Kaiser for her help with the human tissue analysis and Drs. Nagendra Raj and Ranjan Dutta for providing human tissue and cDNA samples.

## Author Contributions

BCS: Conceptualization, data curation and analysis, funding acquisition, writing – original draft, review, and editing. JLW: Conceptualization, data curation, funding acquisition, writing – review and editing.

## Abbreviations

MS: multiple sclerosis
CNS: central nervous system
RRMS: relapsing-remitting multiple sclerosis
SPMS: secondary progressive multiple sclerosis
PPMS: primary progressive multiple sclerosis
PIRA: progression in the absence of relapse activity
IFNγ: interferon γ
EAE: experimental autoimmune encephalomyelitis
LMP7: low molecular weight protein 7
LMP2: low molecular weight protein 2
MECL-1: multicatalytic endopeptidase complex subunit L1
TREM2: triggering receptor expressed on myeloid cells 2
NMD: nonsense mediated decay
P-bodies: processing bodies WML: white matter lesion
NAWM: normal appearing white matter
*i2R-PSMB8*: intron 2 retained *PSMB8*
FL-PSMB8: full-length (canonical) *PSMB8*
*qRT-PCR*: quantitative reverse transcriptase polymerase chain reaction
TBE: tris-borate-ethylenediaminetetraacetic acid buffer
TBS-T: tris-buffered saline
UPF1: upstream frameshift protein 1
pUPF1: phosphorylated upstream frameshift protein 1
HRP: horseradish peroxidase
DMSO: dimethylsulfoxide
CHX: cycloheximide
PBS: phosphate-buffered saline
EDC4: enhancer of mRNA decapping 4
GFP: green fluorescent protein
MHCII: major histocompatibility complex class II
MBP: myelin basic protein
GFAP: glial fibrillary acidic protein

## References

Adinolfi, A., Di Sante, G., Rivignani Vaccari, L., Tredicine, M., Ria, F., Bonvissuto, D., Corvino, V., Sette, C. & Geloso, M. C. 2023. Regionally restricted modulation of Sam68 expression and Arhgef9 alternative splicing in the hippocampus of a murine model of multiple sclerosis. Frontiers in Molecular Neuroscience, 15.

Behm-Ansmant, I. & Izaurralde, E. 2006. Quality control of gene expression: a stepwise assembly pathway for the surveillance complex that triggers nonsense-mediated mrna decay. Genes & Development, 20, 391–398.

Brosnan, C. F. & Raine, C. S. 2013. The astrocyte in multiple sclerosis revisited. Glia, 61, 453–65.

Chang, W.-L. & Tarn, W.-Y. 2009. A role for transportin in deposition of TTP to cytoplasmic RNA granules and MRNA decay. Nucleic Acids Research, 37, 6600–6612.

Chomyk, A. M., Volsko, C., Tripathi, A., Deckard, S. A., Trapp, B. D., Fox, R. J. & Dutta, R. 2017. Dna methylation in demyelinated multiple sclerosis hippocampus. Scientific Reports, 7, 8696.

Chuang, T.-W., Chang, W.-L., Lee, K.-M. & Tarn, W.-Y. 2013. The Rna-binding protein Y14 inhibits mrna decapping and modulates processing body formation. Molecular Biology of the Cell, 24, 1–13.

Compston, A. & Coles, A. 2008. Multiple sclerosis. The Lancet, 372, 1502–1517.

Confavreux, C., Vukusic, S., Moreau, T. & Adeleine, P. 2000. Relapses and progression of disability in multiple sclerosis. N Engl J Med, 343, 1430–8.

Daniels, B. P., Jujjavarapu, H., Durrant, D. M., Williams, J. L., Green, R. R., White, J. P., Lazear, H. M., Gale, M., Jr., Diamond, M. S. & Klein, R. S. 2017. Regional astrocyte Ifn signaling restricts pathogenesis during neurotropic viral infection. J Clin Invest, 127, 843–856.

De Rossi, P., Buggia-Prévot, V., Clayton, B. L. L., Vasquez, J. B., Van Sanford, C., Andrew, R. J., Lesnick, R., Botté, A., Deyts, C., Salem, S., Rao, E., Rice, R. C., Parent, A., Kar, S., Popko, B., Pytel, P., Estus, S. & Thinakaran, G. 2016. Predominant expression of Alzheimer’s disease-associated BIN1 in mature oligodendrocytes and localization to white matter tracts. Molecular Neurodegeneration, 11, 59.

Di Stefano, B., Luo, E.-C., Haggerty, C., Aigner, S., Charlton, J., Brumbaugh, J., Ji, F., Rabano Jiménez, I., Clowers, K. J., Huebner, A. J., Clement, K., Lipchina, I., De Kort, M. A. C., Anselmo, A., Pulice, J., Gerli, M. F. M., Gu, H., Gygi, S. P., Sadreyev, R. I., Meissner, A., Yeo, G. W. & Hochedlinger, K. 2019. The Rna Helicase DDX6 Controls Cellular Plasticity by Modulating P-Body Homeostasis. Cell Stem Cell, 25, 622–638.e13.

Dutta, R., Mcdonough, J., Yin, X., Peterson, J., Chang, A., Torres, T., Gudz, T., Macklin, W. B., Lewis, D. A., Fox, R. J., Rudick, R., Mirnics, K. & Trapp, B. D. 2006. Mitochondrial dysfunction as a cause of axonal degeneration in multiple sclerosis patients. Annals of Neurology, 59, 478–489.

Ebstein, F., Kloetzel, P. M., Kruger, E. & Seifert, U. 2012. Emerging roles of immunoproteasomes beyond Mhc class I antigen processing. Cell Mol Life Sci, 69, 2543–58.

Feinstein, A., Freeman, J. & Lo, A. C. 2015. Treatment of progressive multiple sclerosis: what works, what does not, and what is needed. Lancet Neurol, 14, 194–207.

Griffin, T. A., Nandi, D., Cruz, M., Fehling, H. J., Kaer, L. V., Monaco, J. J. & Colbert, R. A. 1998. Immunoproteasome Assembly: Cooperative Incorporation of Interferon γ (Ifn-γ)–inducible Subunits. Journal of Experimental Medicine, 187, 97–104.

Heink, S., Ludwig, D., Kloetzel, P. M. & Kruger, E. 2005. Ifn-γ-induced immune adaptation of the proteasome system is an accelerated and transient response. Proceedings of the National Academy of Sciences, 102, 9241–9246.

Hurwitz, B. J. 2009. The diagnosis of multiple sclerosis and the clinical subtypes. Ann Indian Acad Neurol, 12, 226–30.

Janssen, W. J., Danhorn, T., Harris, C., Mould, K. J., Lee, F. F., Hedin, B. R., D’alessandro, A., Leach, S. M. & Alper, S. 2020. Inflammation-Induced Alternative Pre-mrna Splicing in Mouse Alveolar Macrophages. G3 (Bethesda), 10, 555–567.

Jiang, L., Wang, M., Lin, S., Jian, R., Li, X., Chan, J., Dong, G., Fang, H., Robinson, A. E., Aguet, F., Anand, S., Ardlie, K. G., Gabriel, S., Getz, G., Graubert, A., Hadley, K., Handsaker, R. E., Huang, K. H., Kashin, S., Macarthur, D. G., Meier, S. R., Nedzel, J. L., Nguyen, D. Y., Segrè, A. V., Todres, E., Balliu, B., Barbeira, A. N., Battle, A., Bonazzola, R., Brown, A., Brown, C. D., Castel, S. E., Conrad, D., Cotter, D. J., Cox, N., Das, S., De Goede, O. M., Dermitzakis, E. T., Engelhardt, B. E., Eskin, E., Eulalio, T. Y., Ferraro, N. M., Flynn, E., Fresard, L., Gamazon, E. R., Garrido-Martín, D., Gay, N. R., Guigó, R., Hamel, A. R., He, Y., Hoffman, P. J., Hormozdiari, F., Hou, L., Im, H. K., Jo, B., Kasela, S., Kellis, M., Kim-Hellmuth, S., Kwong, A., Lappalainen, T., Li, X., Liang, Y., Mangul, S., Mohammadi, P., Montgomery, S. B., Muñoz-Aguirre, M., Nachun, D. C., Nobel, A. B., Oliva, M., Park, Y., Park, Y., Parsana, P., Reverter, F., Rouhana, J. M., Sabatti, C., Saha, A., Skol, A. D., Stephens, M., Stranger, B. E., Strober, B. J., Teran, N. A., Viñuela, A., Wang, G., Wen, X., Wright, F., Wucher, V., Zou, Y., Ferreira, P. G., Li, G., Melé, M., Yeger-Lotem, E., Barcus, M. E., Bradbury, D., Krubit, T., Mclean, J. A., Qi, L., Robinson, K., Roche, N. V., Smith, A. M., Sobin, L., et al. 2020. A Quantitative Proteome Map of the Human Body. Cell, 183, 269–283.e19.

Kashima, I., Yamashita, A., Izumi, N., Kataoka, N., Morishita, R., Hoshino, S., Ohno, M., Dreyfuss, G. & Ohno, S. 2006. Binding of a novel Smg-1–Upf1–eRF1–eRF3 complex (Surf) to the exon junction complex triggers Upf1 phosphorylation and nonsense-mediated mrna decay. Genes & Development, 20, 355–367.

Kiianitsa, K., Kurtz, I., Beeman, N., Matsushita, M., Chien, W.-M., Raskind, W. H. & Korvatska, O. 2021. Novel TREM2 splicing isoform that lacks the V-set immunoglobulin domain is abundant in the human brain. Journal of Leukocyte Biology, 110, 829–837.

Lee, H.-G., Lee, J.-H., Flausino, L. E. & Quintana, F. J. 2023. Neuroinflammation: An astrocyte perspective. Science Translational Medicine, 15, eadi7828.

Lee, J., Villarreal, O. D., Chen, X., Zandee, S., Young, Y. K., Torok, C., Lamarche-Vane, N., Prat, A., Rivest, S., Gosselin, D. & Richard, S. 2020. Quaking Regulates Microexon Alternative Splicing of the Rho GTPase Pathway and Controls Microglia Homeostasis. Cell Reports, 33, 108560.

Marchese, E., Valentini, M., Di Sante, G., Cesari, E., Adinolfi, A., Corvino, V., Ria, F., Sette, C. & Geloso, M. C. 2021. Alternative splicing of neurexins 1–3 is modulated by neuroinflammation in the prefrontal cortex of a murine model of multiple sclerosis. Experimental Neurology, 335, 113497.

Miller, D. H. & Leary, S. M. 2007. Primary-progressive multiple sclerosis. Lancet Neurol, 6, 903–12.

Müller, J., Cagol, A., Lorscheider, J., Tsagkas, C., Benkert, P., Yaldizli, Ö., Kuhle, J., Derfuss, T., Sormani, M. P., Thompson, A., Granziera, C. & Kappos, L. 2023. Harmonizing Definitions for Progression Independent of Relapse Activity in Multiple Sclerosis: A Systematic Review. Jama Neurology, 80, 1232–1245.

Orre, M., Kamphuis, W., Dooves, S., Kooijman, L., Chan, E. T., Kirk, C. J., Dimayuga Smith, V., Koot, S., Mamber, C., Jansen, A. H., Ovaa, H. & Hol, E. M. 2013. Reactive glia show increased immunoproteasome activity in Alzheimer’s disease. Brain, 136, 1415–31.

Paraboschi, E. M., Rimoldi, V., Soldà, G., Tabaglio, T., Dall’osso, C., Saba, E., Vigliano, M., Salviati, A., Leone, M., Benedetti, M. D., Fornasari, D., Saarela, J., De Jager, P. L., Patsopoulos, N. A., D’alfonso, S., Gemmati, D., Duga, S. & Asselta, R. 2014. Functional variations modulating Prkca expression and alternative splicing predispose to multiple sclerosis. Human Molecular Genetics, 23, 6746–6761.

Putscher, E., Hecker, M., Fitzner, B., Boxberger, N., Schwartz, M., Koczan, D., Lorenz, P. & Zettl, U. K. 2022. Genetic risk variants for multiple sclerosis are linked to differences in alternative pre-mrna splicing. Frontiers in Immunology, 13.

Robinson, E. K., Jagannatha, P., Covarrubias, S., Cattle, M., Smaliy, V., Safavi, R., Shapleigh, B., Abu-Shumays, R., Jain, M., Cloonan, S. M., Akeson, M., Brooks, A. N. & Carpenter, S. 2021. Inflammation drives alternative first exon usage to regulate immune genes including a novel iron-regulated isoform of Aim2. Elife, 10.

Shaw, B. C., Snider, H. C., Turner, A. K., Zajac, D. J., Simpson, J. F. & Estus, S. 2022. An Alternatively Spliced TREM2 Isoform Lacking the Ligand Binding Domain is Expressed in Human Brain. Journal of Alzheimer’s Disease, 87, 1647–1657.

Smith, B. C., Sinyuk, M., Jenkins, J. E., Psenicka, M. W. & Williams, J. L. 2020. The impact of regional astrocyte interferon-γ signaling during chronic autoimmunity: a novel role for the immunoproteasome. Journal of Neuroinflammation, 17, 184.

Teixiera, D., Sheth, U., Valencia-Sanchez, M. A., Brengues, M. & Parker, R. 2005. Processing bodies require Rna for assembly and contain nontranslating mRNAs. Rna, 11, 371–382.

Thompson, A. J., Kermode, A. G., Wicks, D., Macmanus, D. G., Kendall, B. E., Kingsley, D. P. & Mcdonald, W. I. 1991. Major differences in the dynamics of primary and secondary progressive multiple sclerosis. Ann Neurol, 29, 53–62.

Trapp, B. D. & Nave, K. A. 2008. Multiple sclerosis: an immune or neurodegenerative disorder? Annu Rev Neurosci, 31, 247–69.

Werkman, I., Sikkema, A. H., Versluijs, J. B., Qin, J., De Boer, P. & Baron, W. 2020. TLR3 agonists induce fibronectin aggregation by activated astrocytes: a role of pro-inflammatory cytokines and fibronectin splice variants. Scientific Reports, 10, 532.

Williams, J. L., Manivasagam, S., Smith, B. C., Sim, J., Vollmer, L. L., Daniels, B. P., Russell, J. H. & Klein, R. S. 2020. Astrocyte-T cell crosstalk regulates region-specific neuroinflammation. Glia, 68, 1361–1374.

Wingerchuk, D. M. & Carter, J. L. 2014. Multiple sclerosis: current and emerging disease-modifying therapies and treatment strategies. Mayo Clin Proc, 89, 225–40.

Zajac, D. J., Simpson, J., Zhang, E., Parikh, I. & Estus, S. 2023. Expression of INPP5d Isoforms in Human Brain: Impact of Alzheimer’s Disease Neuropathology and Genetics. Genes, 14, 763.

Zheng, J. & Bizzozero, O. A. 2011. Decreased activity of the 20s proteasome in the brain white matter and gray matter of patients with multiple sclerosis. J Neurochem, 117, 143–53.

