## Supplemental Figure 1, Supplemental Figure 2, Supplemental Figure 3 for "A novel *PSMB8* isoform associated with multiple sclerosis lesions induces P-body formation"

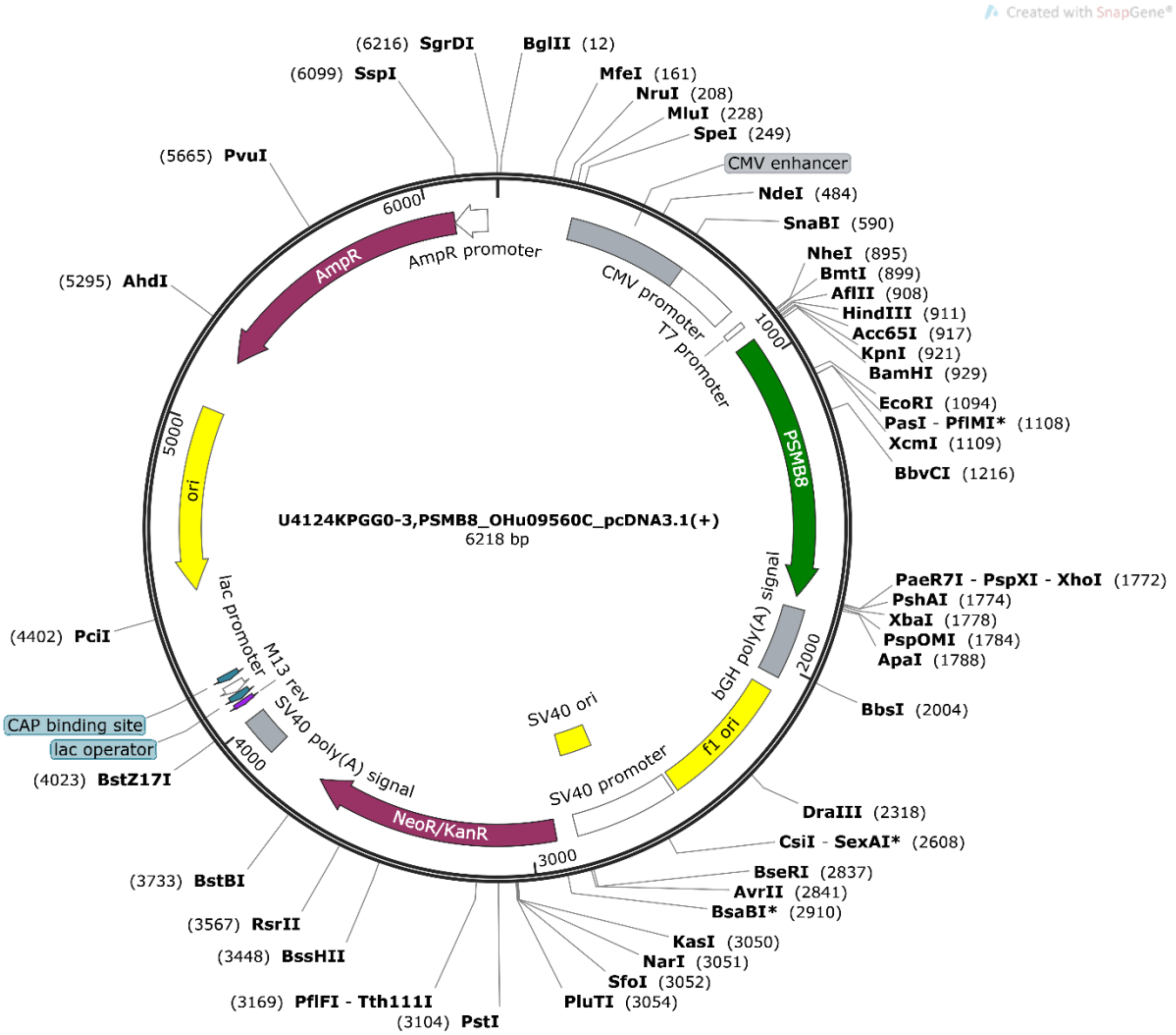


**Supplemental Figure 1.** Vector map of *FL-PSMB8* plasmid used in transfections. This plasmid was purchased from Genscript, catalog number SC1200, GenEZ ORF Clone: PSMB8_OHu09560C_pcDNA3.1(+). This is the canonical exon 1B containing *PSMB8*, NM_148919.4.

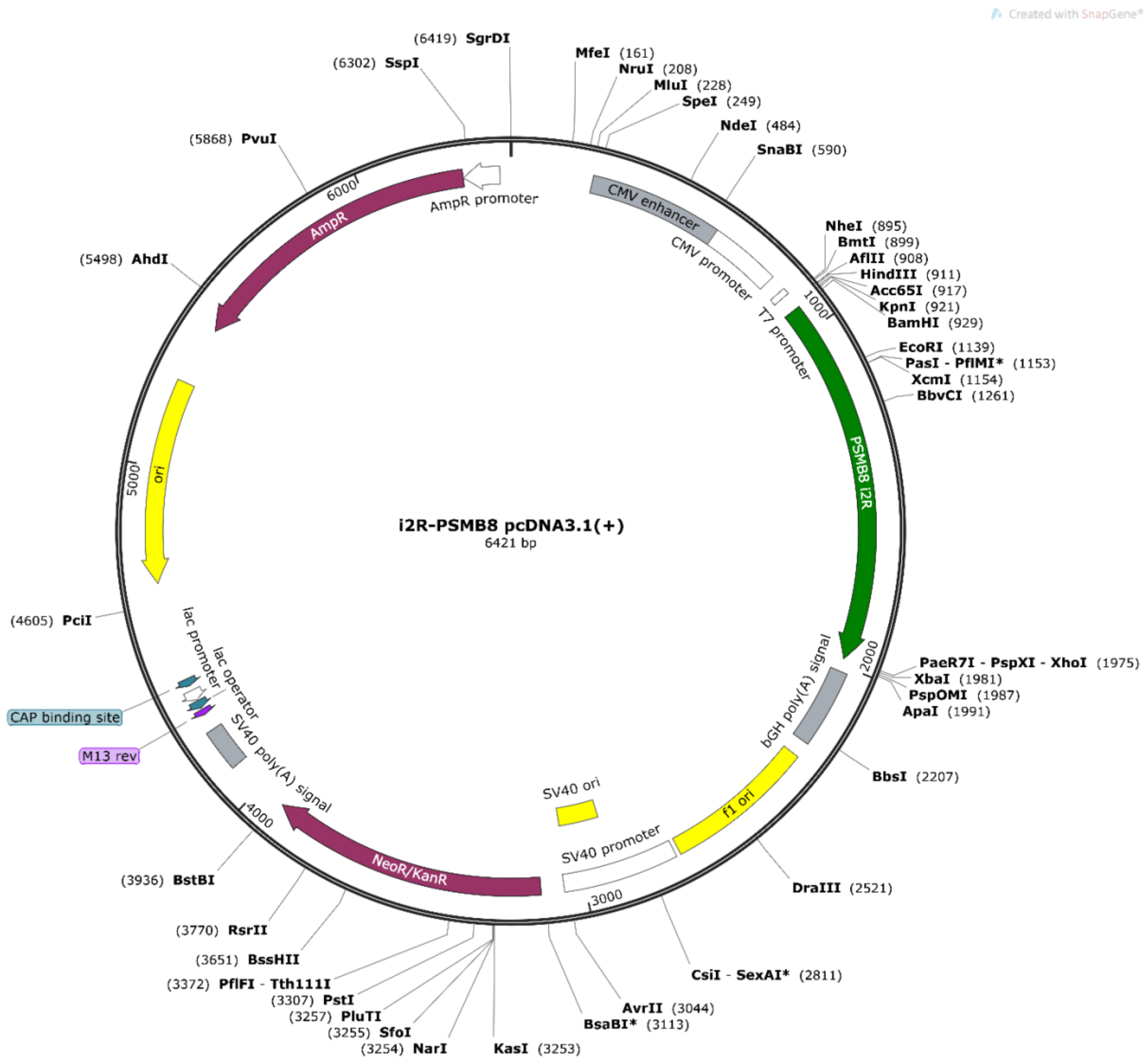


**Supplemental Figure 2.** Vector map of *i2R-PSMB8*. This was a custom gene synthesis from Genscript cloned into pcDNA3.1. This is identical cDNA as the exon 1B containing *PSMB8* but with intron 2 retained.

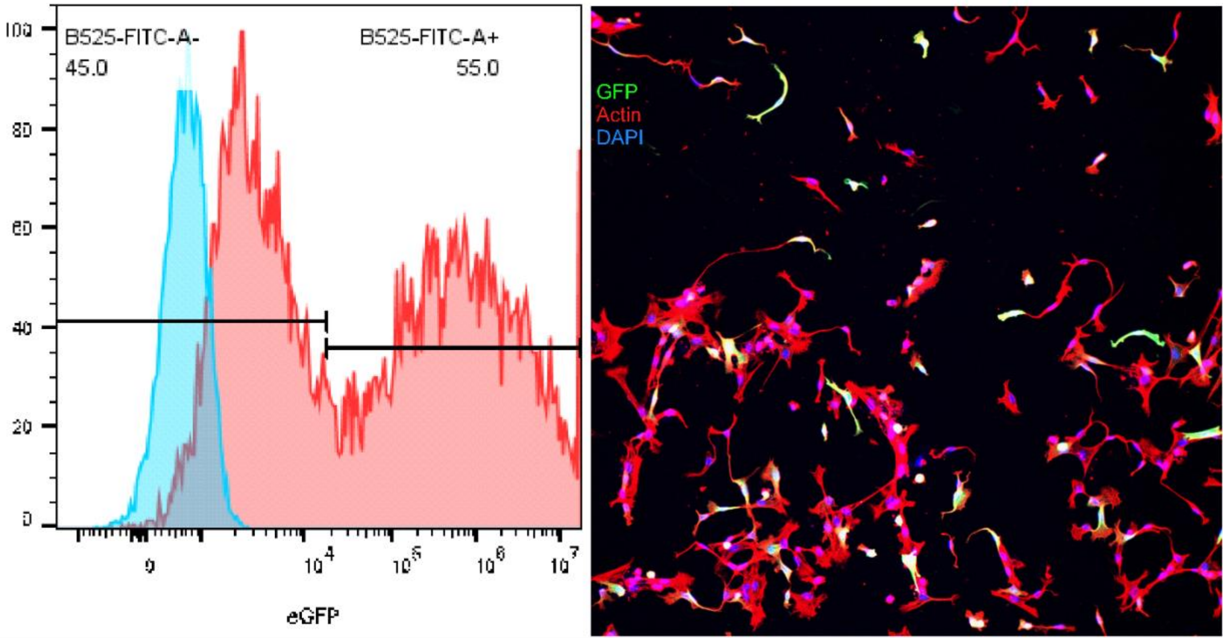


**Supplemental Figure 3.** Transfection efficiency of primary human cortical astrocytes. Left: flow cytometry of GFP-transfected primary human cortical astrocytes (red) or non-transfected controls (blue). Right: fluorescence microscopy image of GFP-transfected primary human cortical astrocytes. Actin is shown in red and DAPI in blue to mark all cells, and GFP, shown in green, as an indicator of transfection. Astrocytes sourced from ScienCell and transfections performed using a Lonza Nucleofector 4 device with 6 µg DNA per $10^6$ cells under program DR114.
